# Effects of I*_h_* and TASK-like shunting current on dendritic impedance in layer 5 pyramidal-tract neurons

**DOI:** 10.1101/2021.01.08.425962

**Authors:** Craig Kelley, Salvador Dura-Bernal, Samuel A. Neymotin, Srdjan D. Antic, Nicholas T. Carnevale, Michele Migliore, William W Lytton

## Abstract

Pyramidal neurons in neocortex have complex input-output relationships that depend on their morphologies, ion channel distributions, and the nature of their inputs, but which cannot be replicated by simple integrate-and-fire models. The impedance properties of their dendritic arbors, such as resonance and phase shift, shape neuronal responses to synaptic inputs and provide intraneuronal functional maps reflecting their intrinsic dynamics and excitability. Experimental studies of dendritic impedance have shown that neocortical pyramidal tract neurons exhibit distance-dependent changes in resonance and impedance phase with respect to the soma. We therefore investigated how well several biophysically-detailed multi-compartment models of neocortical layer 5 pyramidal tract neurons reproduce the location-dependent impedance profiles observed experimentally. Each model tested here exhibited location-dependent impedance profiles, but most captured either the observed impedance amplitude or phase, not both. The only model that captured features from both incorporates HCN channels and a shunting current, like that produced by Twik-related acid-sensitive K^+^(TASK) channels. TASK-like channel activity in this model was dependent on local peak HCN channel conductance (*I*_h_). We found that while this shunting current alone is insufficient to produce resonance or realistic phase response, it modulates all features of dendritic impedance, including resonance frequencies, resonance strength, synchronous frequencies, and total inductive phase. We also explored how the interaction of *I*_h_ and a TASK-like shunting current shape synaptic potentials and produce degeneracy in dendritic impedance profiles, wherein different combinations of *I*_h_ and shunting current can produce the same impedance profile.

**New & Noteworthy:** We simulated chirp current stimulation in the apical dendrites of 5 biophysically-detailed multi-compartment models of neocortical pyramidal tract neurons and found that a combination of HCN channels and TASK-like channels produced the best fit to experimental measurements of dendritic impedance. We then explored how HCN and TASK-like channels can shape the dendritic impedance as well as the voltage response to synaptic currents.

## Introduction

The pyramidal cells (PCs) found in layer 5 (L5) of neocortex generate the main outputs of cortical circuits: spike trains propagating along axons that project to various cortical and subcortical structures, exerting top down control over other brain areas and motor function [3, 21, 22, 40, 50, 77]. In order to produce their outputs, L5 PCs integrate inputs from other cortical layers, other cortical areas, and thalamus [2, 43, 47, 55, 59, 80]. There is great diversity among PCs in L5, not just in their morphologies and projections, but also in their spiking activity, with some PCs having high spontaneous firing rates while others’ firing rates are closely correlated with the activity of neurons in the surrounding population [43]. The balance of excitatory and inhibitory inputs and the electrotonic structure of PCs are key in understanding how they generate their outputs and exert top-down control over other parts of the nervous system.

In this study we focused on pyramidal tract neurons (PTs; also called thick-tufted cells), one of the 3 major classes of cortical PCs. **1**. PTs project to subcortical structures and include corticospinal, corticobulbar, and corticopontine cells as well as projections to the medullary pyramids [13, 21]. They also send collateral projections to thalamus. **2**. Intratelencephalic neurons (ITs), also called thin-tufted or commissural cells, include corticostriatal and corticocortical cells and project to other cortical areas [56]. **3**. Corticothalamic neurons (CTs) project to ipsilateral thalamus [77]. A major physiological factor distinguishing PTs from ITs and CTs is the high expression of the hyperpolarization-activated cyclic nucleotide–gated (HCN) channel, a nonselective voltage-gated cation channel responsible for the h-current (I_*h*_) [13, 56, 68]. High expression of HCN channels profoundly affects the subthreshold filtering properties of neuronal membrane.

The electrical properties of the passive neuronal membrane are very similar to those of a parallel RC circuit, with the response of membrane potential to currents dropping off at frequencies above the “natural frequency” at 1/2*π*RC Hz (low-pass filtering). Under the right circumstances, however, voltage-gated ion channels can produce a “phenomenological inductance” [7, 8] that can, like a physical inductor in an RLC circuit, generate resonance: an enhanced voltage response over an intermediate range of frequencies [45, 46]. Phenomenological inductance is most likely to be seen when channels with slow gating are present, such as HCN channels and delayed rectifier K channels [29, 58, 71]. Resonance becomes apparent when currents through these channels are prominent enough and lag sufficiently far behind fluctuations of membrane potential [30].

The filtering properties of the neuronal membrane have been characterized as impedance profiles measured at subthreshold voltages [10, 12, 57, 58]. A common experimental method for probing neuronal impedance is to stimulate the neuron by injecting a chirp current waveform: a constant-amplitude, sinusoidal waveform whose instantaneous frequency increases from low to high over time [12, 57]. In this study, we use a linear chirp stimulus whose instantaneous frequency increases linearly from 0.5-20 Hz over 20s [13, 75]. Impedance amplitude (|*Z*|) characterizes voltage response with respect to stimulus frequency. The resonant peak (resonant frequency, f_*res*_) is found at the frequency where the constant amplitude current stimulus causes the greatest peak-to-peak changes in membrane potential. *I*_h_-mediated resonance has been observed in a wide variety of species and neuronal cell types [30, 75, 82], and is proposed to impart neurons with the ability to discriminate inputs by frequency [4, 12, 29]. In addition to responding more strongly at certain frequencies (resonance), *I*_h_ also provides another property characteristic of inductive circuits: a shift of response phase (Φ). Given a sinusoidal current stimulus, the peaks of V_memb_ may occur before (lead), after (lag), or synchronous with peaks in the stimulating current [52]. The frequency at which a peaks in the stimulating current and peaks in V_memb_ is referred to as the synchronous frequency [13]. The phenomenological inductance produced by *I*_h_ opposes capacitive delay imparted by the neuronal membrane and produces phase lead at some frequencies. *I*_h_ has thus been proposed as a mechanism for compensating location-dependent capacitive delays of dendritic inputs seen at the soma, ensuring that simultaneous synaptic inputs dispersed across the dendritic arbor are coincident in the soma [76].

To illustrate these effects, we modified the standard passive neuronal model by adding an inductive circuit which mimics some of the properties of *I*_h_ (Fig. 1A). The resistor (R) stands in for the conductance of *I*_h_; the battery (E), its reversal potential; and the inductance (L), the phenomenological inductance it generates. We used an extremely high inductance of 10 kH to show obvious effects. Adding the inductive circuit changes the low-pass filter properties of the passive neuron (Fig. 1B, dashed lines) to those of a resonator (solid lines). The inductance also increases impedance phase, even creating phase lead at low frequencies, where the impedance phase profile (ZPP) is greater than zero (Fig. 1C). The inductance that shapes the impedance amplitude profile (ZAP) and ZPP also influences synaptic potentials In dendrites equipped with inductance, the EPSP becomes faster (peak occurs sooner) and narrower (half width decreases) Fig. 1D. Resonance in the ZAP is associated with narrowing the shape of the EPSP, which is consistent with the effects of HCN channels in dendrites [34, 81]. Higher impedance phase in the ZPP is associated with earlier peak V_memb_ in the EPSP, even showing synaptic phase lead with peak somatic V_memb_ preceding peak synaptic current in the dendrite. While the phenomenological inductance produced by HCN channels is not sufficient for phase lead in synaptic potentials, we will see that increased impedance phase compensates for membrane capacitance and reduces the delay between peak synaptic conductance in the dendrites and peak V_memb_ at the soma.

**Fig 1.**
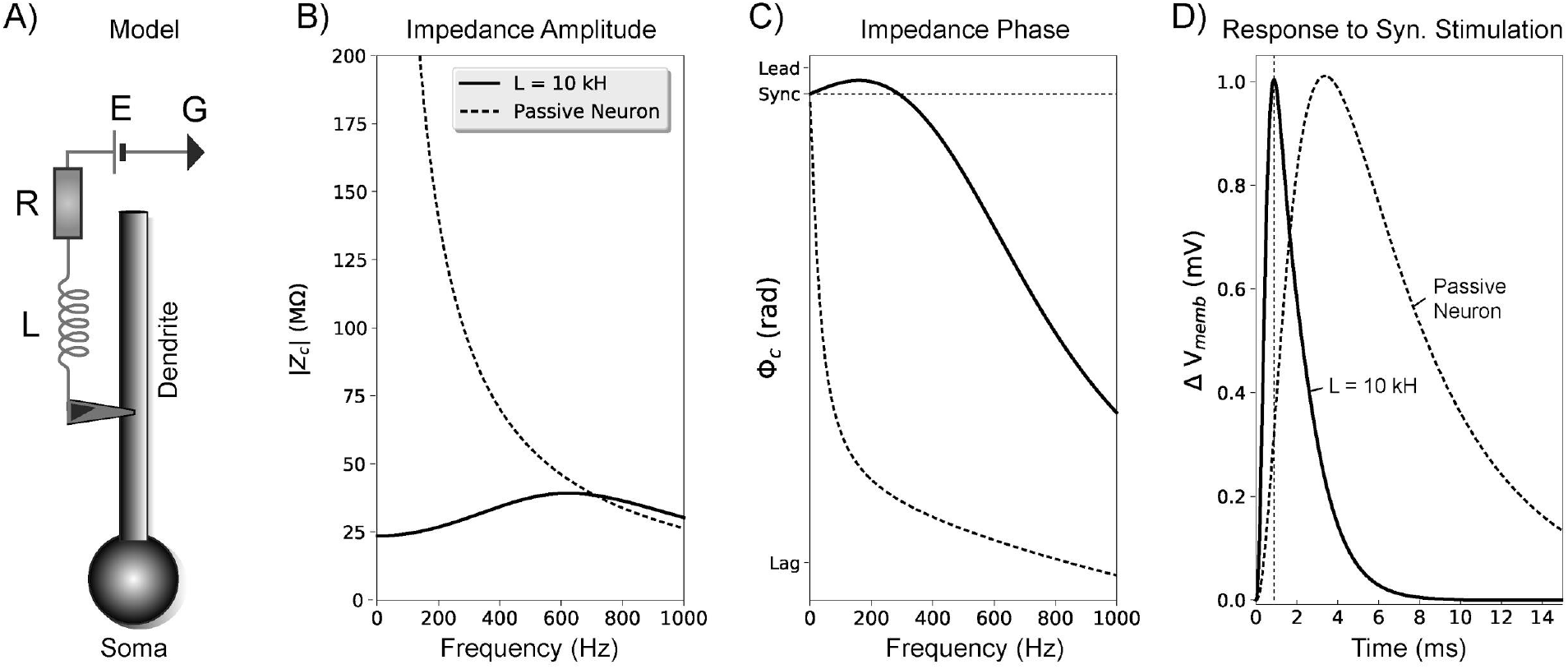
Inductance influences neuronal impedance and the response to synaptic stimulation. (A) A simple, passive neuron model (soma and dendrite with membrane capacitance) was connected to a series circuit with an inductor (L = 10 kH), resistor (R = 25 MΩ), and battery (E = −70 mV) to illustrate some of the effects of inductance on impedance and synaptic potentials. We computed impedance between the center of the dendrite and the center of the soma with this circuit attached (solid black lines) and without it (dashed black lines). (B) The inductive circuit combined with membrane capacitance from the neuron produces resonance. In the passive neuron alone, impedance amplitude falls off with frequency. (C) The inductive circuit also increases impedance phase across all frequencies, with positive inductive/leading phase (voltage peak precedes current peak for an oscillatory input) seen at low frequencies. The horizontal dotted line indicates 0 radian phase shift between the stimulating current and voltage response at the soma (i.e. synchrony). (D) Effects of increased inductance on EPSPs measured at the soma: peak voltage is earlier due to inductive phase, and the waveform is narrower due to resonance. Time of peak synaptic conductance is indicated by the vertical dotted line.

I_*h*_ has other dramatic effects on the intrinsic dynamics and excitability of neurons. It acts as a pacemaker current, supporting regular- and burst-firing modes [64]. It mediates the sag potential observed during hyperpolarization and spike-frequency adaptation during suprathreshold depolarization [56, 64]. I_*h*_ supports coincidence detection, affects temporal summation [11, 14, 39], and has been suggested to determine the frequency response of neuronal membrane potential (V_memb_) in response to weak alternating electric fields, like that produced by transcranial current stimulation [73]. Additionally, HCN channels have been shown to have paradoxical effects on excitatory post-synaptic potentials (EPSPs), enhancing spiking in response to EPSPs when the spike threshold is low and inhibiting spiking in response to EPSPs when the spike threshold is high [19]. Recent modeling studies have suggested that this dual role could be attributed to interactions between HCN channels and a shunting current, most likely that produced by Twik-related acid-sensitive K^+^(TASK) channels [16, 48].

The relatively high expression of HCN in PTs endows them with resonance, giving the properties of a band-pass filter [13, 30, 75]. We here report that five previously developed, biophysically-detailed multi-compartment models of neocortical PTs exhibit dendrite-location-dependent impedance profiles with resonant frequencies and synchronous frequencies increasing with distance from the soma [16, 18, 23, 38, 54]. Four of the five models have resonant frequencies in line with experimental findings, ranging from 4-9 Hz [13, 75], while the fifth produced resonant frequencies above this range. Two of the five models have synchronous frequencies in line with experimental data, ranging from 3.5-7 Hz [13], while the other three produced synchronous frequencies below this range. Only one PT model, which includes both *I*_h_ and a TASK-like shunting current, produced realistic impedance amplitude and phase profiles. We added TASK-like channels to one of the PT models that originally only produced resonant frequencies matching experimental findings. This addition produced realistic impedance amplitude and phase profiles with resonant and synchronous frequencies withing the experimental range. We also examined how *I*_h_ and the TASK-like shunting current interact to produce and modulate dendritic resonance, inductive phase, and the properties of EPSPs.

## Methods

The biophysically-detailed models studied here were developed for and published in previous studies. [1, 16, 23, 38, 54] All simulations presented here were performed using NEURON version 7.8.0 [25, 26]. The code developed for simulation, data analysis, and visualization was written in Python, and it is available on [GitHub](https://github.com/suny-downstate-medical-center/L5PYR_Resonance) and [ModelDB](https://senselab.med.yale.edu/modeldb/ShowModel?model=266851).

### Models

The simplified neuron model presented in Fig. 1 had a single-compartment, spherical soma with radius 5 *µ*m, and a single three-compartment dendrite 75 *µ*m long and 10 *µ*m in diameter. All compartments had a membrane capacitance of 1 *µ*F/cm^2^, passive conductance 0.2 mS/cm^2^, and passive reversal potential of −70 mV. To demonstrate the effects of inductance on impedance and V_memb_ dynamics, the cell was connected to an inductor (L = 10 kH), resistor (R = 25 MΩ), and a battery (E = −70 mV) placed in series and connected to ground (Fig. 1A).

We focused our study on 5 biophysically-detailed, multi-compartment models: three models of rat PTs and two models of mouse PTs (Table 1). **Model 1** is based on data from neocortex of Wistar rats, postnatal day (P) 36 [23]. The model was fit to perisomatic and backpropagating spiking activity. Dendritic channels were uniformly distributed with the exceptions of HCN channels and high- and low-voltage activated Ca^2^+ channels. I_*h*_ was uniform in the basal dendrites, while in the apical dendrites I_*h*_ channels were distributed using a density function that increased exponentially with distance from the soma [37, 53]. The density of Ca^2^+ channels was increased near the nexus of the apical tufts forming a “hot-zone” [23]. **Model 2** was based on data from frontal cortex of Sprague-Dawley rats, P21-33, fit using voltage-sensitive dye imaging data with a focus on reproducing dendritic plateau potentials and their propagation toward the soma, dendritic sodium spikelets, and backpropagating action potentials in the basal dendrites [1, 18]. The distribution of I_*h*_ channels was constant in the basal dendrites and increased exponentially with distance from the soma in the apical dendrites. **Model 3** was based on data from somatosensory cortex of Wistar rats [38]. Channel densities were adjusted primarily to account for perisomatic spiking activity, particularly fast action potential repolarization and large amplitude afterhyperpolarization in the axon initial segment. I_*h*_ channels were distributed throughout the dendritic arbor with an exponential increase in density with distance from the soma [37]. It also had M-type K^+^ channels distributed uniformly throughout the dendritic arbor. **Model 4** was based on data from primary motor cortex (M1) of C57Bl/6J mice, P21 [54] . The model was fit based on perisomatic spiking activity and validated by simulating subthreshold somatic resonance. I_*h*_ conductance was constant in the basal dendrites, increased exponentially with distance from the soma along the apical trunk until the nexus with apical dendrite tufts, beyond which the I_*h*_ conductance plateaued at 0.006 S/cm^2^ [20]. **Model 5** was based on **Model 4**; they had identical morphologies [16]. It was modified to include a TASK-like shunting current whose conductivity was coupled to peak I_*h*_ conductivity as described by Migliore & Migliore (2012), along with small changes to fast sodium channel conductance, membrane capacitance, and passive conductance [16]. These changes preserved the perisomatic firing characteristics of the original model and fit experimental data from PT cells in primary motor cortex while also reproducing additional I_*h*_-dependent phenomena observed experimentally [16, 19, 54, 68]. More detailed information regarding the parameters and properties of the models studied here may be found in their original publications [16, 18, 23, 38, 54].

**Table 1.**
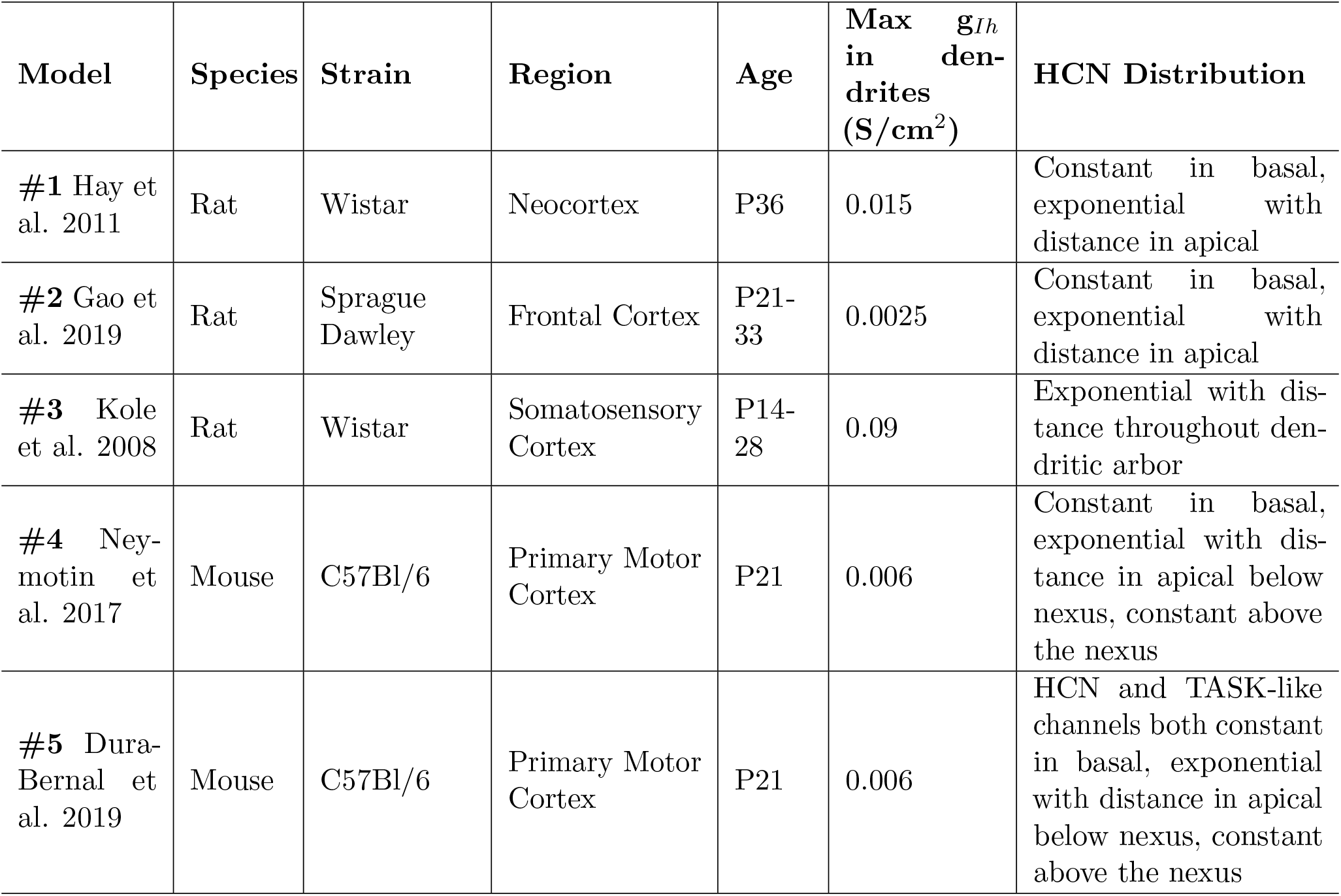
Basic Model Information. Models are specified by either the publication in which they first appeared. Ages are specified by postnatal day age. Under the comments on HCN channel distribution, “exponential with distance” is with respect to the soma.

### Chirp and impedance

We generated impedance profiles for each of these models by stimulating each compartment along the apical trunk with a chirp current waveform and measuring changes in V_*memb*_ at the soma. We used a linear chip stimulus where current (*I*_*in*_) is defined as:

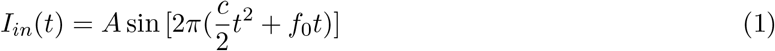

where *c* = (*f*_1_-*f*_0_) / *T* , *f*_0_ is the initial frequency, *f*_1_ is the final frequency, and T is the duration of the frequency sweep. *A*, the stimulus amplitude, was chosen such that excursions in V_memb_ about V_*rest*_ were symmetrical to within 0.01 mV. The instantaneous frequency of *I*_*in*_(*t*) increases linearly with time. When computing impedance in the biophysically-detailed PT models, we used *f*_0_ = 0.5 Hz, *f*_1_ = 20Hz, and *T* = 20 s. It should be noted that commonly used scientific computing software packages like SciPy and MATLAB’s Signal Processing Toolbox include chirp functions that use cosine rather than sine, and a phase shift of −90 degrees must be used to ensure smooth transitions in V_*memb*_ when using these functions to generate stimuli appropriate for impedance analysis [44, 78].

We focused specifically on the transfer impedance between the stimulated dendrite and the soma, which was computed as:

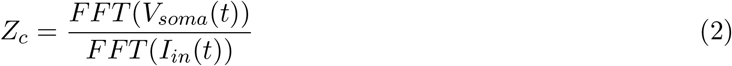

*Z*_*c*_ is a complex valued function where FFT(I_*in*_) is the Fourier transform of the injected current waveform and FFT(V_*m*_) is the Fourier transform of the change in membrane potential at the soma. From the impedance we extract the real valued resistance (*R*) and the imaginary valued reactance (*X*). From *R* and *X* we compute the transfer impedance amplitude as a function of input frequency:

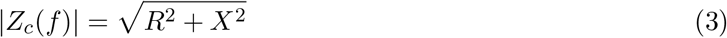

Transfer frequency, *f*_*transfer*_, is defined as the frequency at which |*Z*_*c*_| between the stimulation site and the soma is maximized [13]. In other words, *f*_*transfer*_ is the resonant frequency (*f*_*res*_) of the transfer impedance. Transfer resonance strength (*S*_*c*_) is a dimensionless quantity defined as:

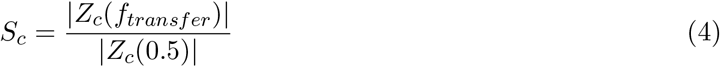

This quantity has been used in previous publications and referred to as “Q factor” or “Q”, but this measure differs entirely the generally accepted definition of Q factor used in the context of resonant electrical circuits [36, 74]. We therefore simply refer to the quantity in Equation (4) as resonance strength.

Transfer impedance phase (Φ_*c*_), which quantifies the temporal relationship between *I*(*t*) and V_memb_ at the soma, is defined as:

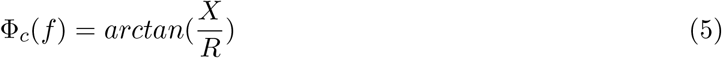

Synchronous frequency between the dendrite and soma is defined as the frequency at whichΦ_*c*_ = 0 and peaks in *I*_*in*_(*t*) and V_*soma*_ are synchronized. WhenΦ_*c*_ *>* 0, the peaks in V_*soma*_ precede *I*(*t*), which is referred to as leading or inductive phase. Total inductive phase [52] is defined as:

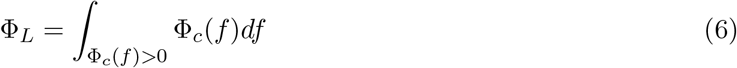

– the area of theΦ_*c*_ curve above zero. If there is no inductive phase andΦ_*c*_ *<* 0 for all frequencies, we set the synchronous frequency to zero.

Because the chirp waveform is not stationary (its instantaneous frequency increases over time), and the discrete Fourier transforms used in Equation 2 to compute impedance assume the signal is stationary, we validated the use of chirp to generate impedance profiles. We compared impedance profiles generated using chirp with impedance profiles generated by stimulating the cell with stationary sinusoidal current waveforms at a single frequency for 5 s, computing the impedance phase and amplitude at that frequency, and repeating for each frequency of interest. We found that impedance amplitudes are nearly identical between the two methods, but there are differences impedance phase (Supplemental Fig. S1 https://doi.org/10.6084/m9.figshare.13322588.v1). For instance, when using chirp to compute impedance in one of the biophysically-detailed models, impedance phase is is practically indistinguishable from 0.5 − 13 Hz using both methods, but phase begins to diverge beyond 13 Hz. We also see in the simplified models that the errors in impedance phase increase at higher frequencies. Since important impedance phase features like synchronous frequency andΦ_*L*_ occur below 13 Hz in PTs, the chirp waveform is suitable for computing impedance phase. However, we recommend caution if one is using chirp to compute impedance phase at higher frequencies. Therefore, when computing impedance for the simple models seen in Fig. 1, we used a 5 s sinusoid at each frequency (0.5 − 1000 Hz in 0.5 Hz increments) rather than chirp.

### Simulations

We ran over 4000 single-cell simulations during the course of this study. For simulating chirp stimulation of the biophysically-detailed PT models, 1 second of simulation-time took roughly 40 seconds of clock-time in NEURON on a Linux system using 2.40 GHz quad-core Intel Xeon CPUs. We simulated chirp current stimulation of each compartment along the apical trunks of each PT model and computed the transfer impedance between the stimulated compartment and the soma. By determining the transfer resonance frequencies and synchronous frequencies along the apical trunk, we observed the location-dependence of the impedance profiles in these PT cell models. For comparisons between the models and experimental data, transfer frequency and synchronous frequency observations were extracted from published data [13, 75] using WebPlotDigitizer [65] and pooled together. Since each observation was made from a different neuron, and it is not indicated how far each measurement is relative to the apical trunk length, we normalized all position data to the farthest observation from the soma.

All synaptic stimulation simulations were performed using NEURON’s AlphaSynapse with a time constant of 1 ms to mimic a unitary, excitatory AMPA synapse [25]. Maximal synaptic conductance was chosen to produce a∼1 mV depolarization in somatic V_memb_ in each model or condition for all synaptic stimulation simulations.

## Results

### Impedance profiles of model PT neurons

Since location-dependent gradients in resonance and impedance phase were not investigated previously in PT models [17, 35], we explore how both impedance amplitude and phase change with distance from the soma in morphologically and biophysically detailed PT models. We measured the impedance profiles of 5 biophysically-detailed multi-compartment models of L5 PTs using a set of simulated 20 s subthreshold chirp-waveform current injections with instantaneous frequency of 0.5 − 20 Hz (Fig. 2A). We simulated stimulation with a subthreshold chirp-waveform at various locations along the apical trunk (Fig. 2B). Changes in membrane potential in response to chirp stimuli were recorded from the stimulated compartments (Fig. 2C, E, G) and at the soma (Fig. 2D, F, G). We computed transfer impedance (Z_*c*_) and associated measures from each of the recorded somatic membrane potential waveforms via Equations 2-4 (Fig. 2I-K). In an example PT model, we see location-dependent changes in the impedance profiles with transfer frequencies, resonance strength, total inductive phase, and synchronous frequencies all increasing along the apical trunk with distance from the soma (Fig. 2J, K). The peaks and contours of the ZAP and ZPP shift to the right in frequency with distance from the soma (Fig. 2J, K).

**Fig 2.**
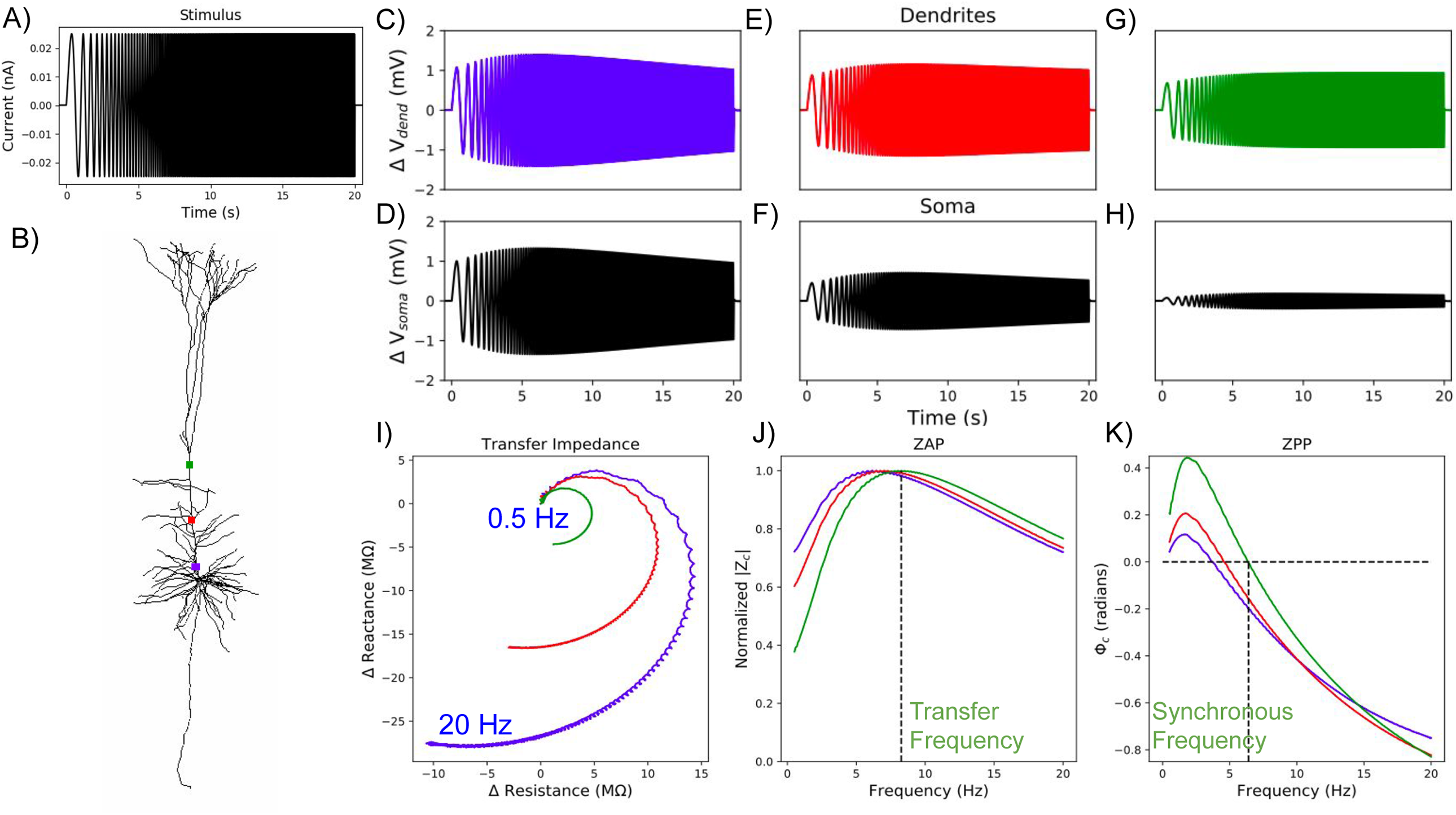
Impedance responses in dendrites of Model 5. (A) Constant amplitude, linear chirp, current waveform which is applied to different points along the apical dendrite. (B) Stimulated locations along the apical trunk: proximal (blue), central (red), and distal(green). We recorded membrane potentials at the stimulated compartments (C, E, G) and at the soma (D, F, H). (I) *Z*_*c*_ was computed from the changes in the membrane potential at the soma and the current stimulus applied to the dendrites. (J) From the transfer impedance amplitude, |*Z*_*c*_|, we compute the transfer resonance frequency, which is indicated by the vertical dashed line for the most distal recording site. (K) From the transfer impedance phase,Φ_*c*_, we compute the synchronous frequency, again indicated by a vertical dashed line for the most distal site.

All of the PT cell models exhibit location-dependent impedance profiles with transfer frequencies and synchronous frequencies increasing with distance from the soma (Fig. 3) [13, 75]. They varied, however, in how well they replicated the full range of experimental data. **Model 3** overestimated the transfer frequency along the apical trunk but exhibited synchronous frequencies within the experimental range. **Models 1, 2, & 4** exhibited realistic transfer frequencies along the apical trunk but underestimated the synchronous frequencies. **Models 1 & 2** even showed no inductive phase, withΦ_*c*_ *<* 0 at all frequencies (5), for large proximal portions of their apical trunks (Fig. 3B). Only **model 5** captured both the transfer and synchronous frequencies observed in experiments.

**Fig 3.**
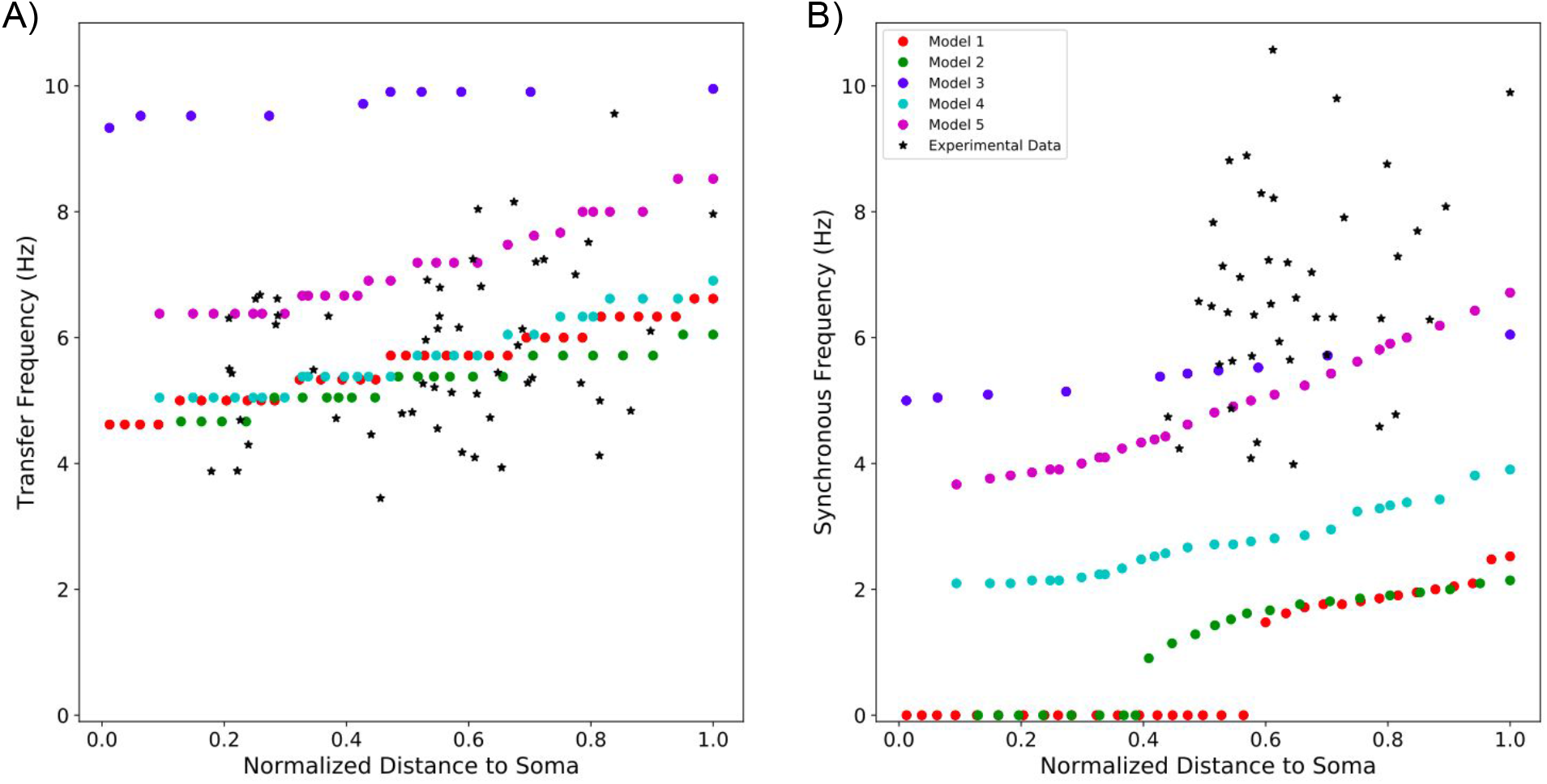
Resonant frequencies and synchronous frequencies of 5 PT models compared to experimental data. (A) Four of the five models show transfer frequencies along the apical trunk within the experimentally observed range. The fifth produced transfer frequencies above this range. Experimental values of transfer frequencies were extracted from Ulrich (2002) and Dembrow et al. (2015). (B) Only two models exhibit synchronous frequencies along the apical trunk which are within the experimental range. The other three models produce synchronous frequencies below this range. Experimental values of synchronous frequencies were extracted from Dembrow et al. (2015)

**Model 5** produced greater total inductive phase along its apical trunk than any of the other models (Fig. 4).Φ_*L*_ (6) between the distal end of the apical trunk and the soma was roughly 7x higher in **model 5** with the both HCN and TASK-like channels compared to its earlier incarnation (see Methods, Table 1) **model 4** (Fig. 4A). As an example, we present ZPPs from the same segment in **models 4 & 5**, roughly half the length of the apical trunk (136.4 *µ*m) from the soma (Fig. 4B). PeakΦ_*c*_ in **model 5** is more than double that in **model 4**, andΦ_*c*_ remains higher in **model 5** than **model 4** for all frequencies probed. The optimal frequency for leading phase remained around 2 Hz in both models however. In the time domain, this means that V_memb_ at the soma leads a 2 Hz sinusoidal stimulating current halfway along the apical trunk by roughly 17 ms in **model 5**, whereas they are practically synchronous in **model 4** (Fig. 4B, inset). Although the increasedΦ_*L*_ is not sufficient to produce phase lead in the EPSP, increasedΦ_*c*_ partially compensates for the capacitive delay in EPSP arrival time at the soma (Fig. 4C). When synaptic stimulation halfway along the apical trunk produces a 1 mV amplitude EPSP in the soma, peak V_memb_ occurs roughly 1 ms sooner in **model 5** than in **model 4**. This difference is consistent across a range of EPSP amplitudes (0.5 − 2 mV, data not shown), and we expect it to remain consistent within the subthreshold range.

**Fig 4.**
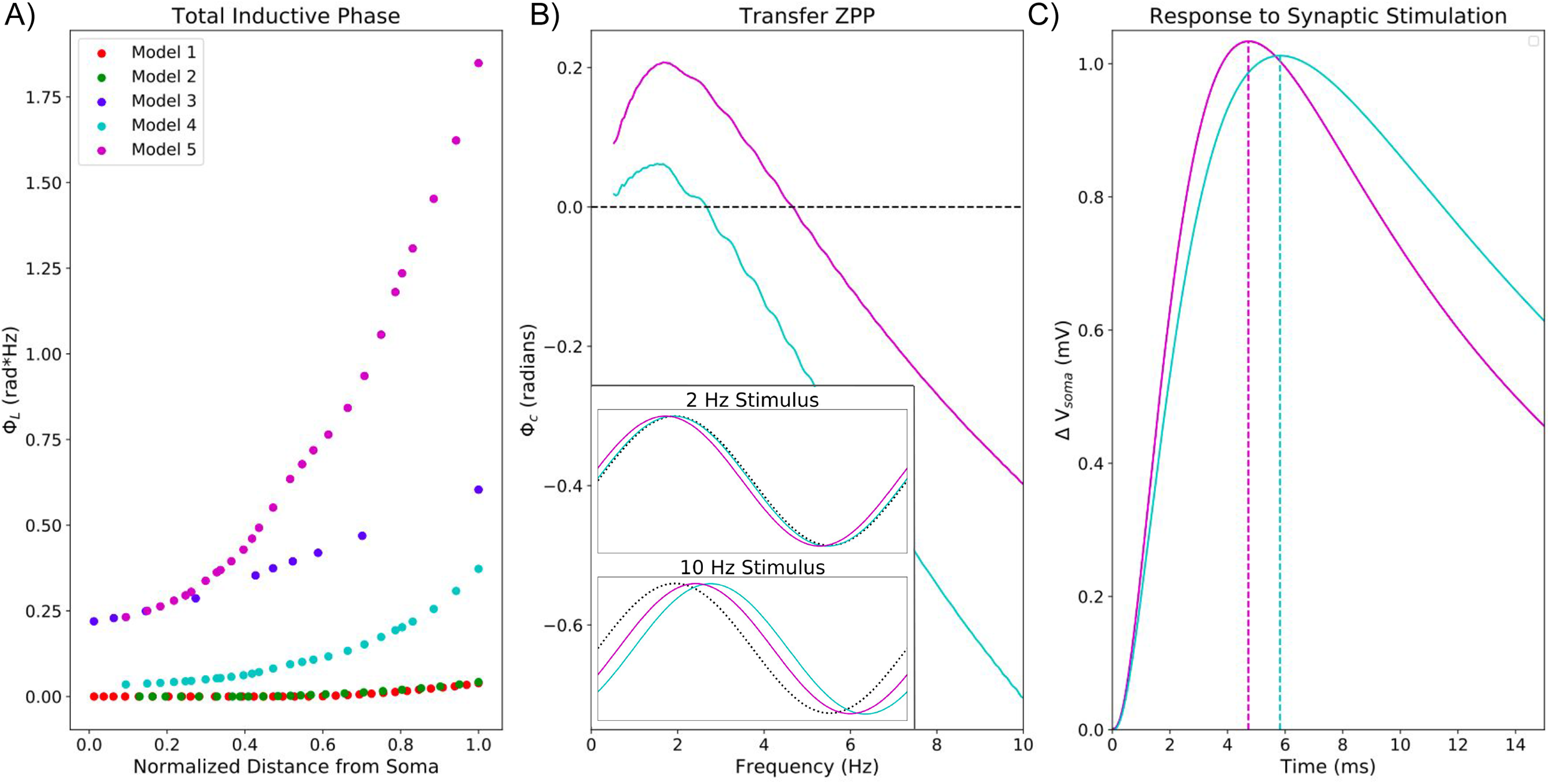
The impedance phase in PT models and its implications for synaptic potentials. (A) **Model 5** exhibits much greater total inductive phase along the apical trunk compared to the other models. (B) Comparison of two models’ ZPPs from halfway along the apical trunk (136.4 *µ*m from the soma) showingΦ_*c*_ is greater in **model 5** than in **model 4** for all frequencies probed. Inset shows somatic V_memb_ response to 2 Hz and 10 Hz sinusoidal stimuli in the time domain from both models. At 2 Hz, V_memb_ leads the stimulating current by roughly 17 ms in **model 5**, while they are nearly synchronous in **model 4**. At 10 Hz, lag in V_memb_ is reduced in **model 5** compared to **model 4**. Dotted black lines indicate the stimulating current waveform. (C) Somatic EPSP in response to synaptic stimulation in both models at the same point along the apical trunk. Peak V_memb_ occurs more than 1ms earlier in **model 5** than in **model 4**.

### I_*h*_, TASK-like shunting current, and dendritic impedance

A combination of I_*h*_ and TASK-like shunting current produced the best approximation of experimentally observed dendritic impedance profiles in PTs (Fig. 5). **Model 5** was the only PT model which included a TASK-like shunting current that was coupled to peak I_*h*_ conductivity [48]. We repeated our simulations on **model 5** with different models of the HCN channel that do not include an additional shunting current in order to determine what produced its biologically realistic impedance profiles. We computed transfer and synchronous frequencies along the apical trunk using models of HCN from Kole et al. (2006) and Harnett et al. (2015). While using the other two HCN models reduced the transfer frequencies along the apical trunk, these remained well within the observed range (Fig. 5A). The different models of HCN had dramatic effects on the phase response however. The Harnett et al. (2015) model reduced synchronous frequency by roughly one half across the apical trunk. The Kole et al. (2006) model produced zero inductive phase along more than half the length of apical trunk (Fig. 5B).

**Fig 5.**
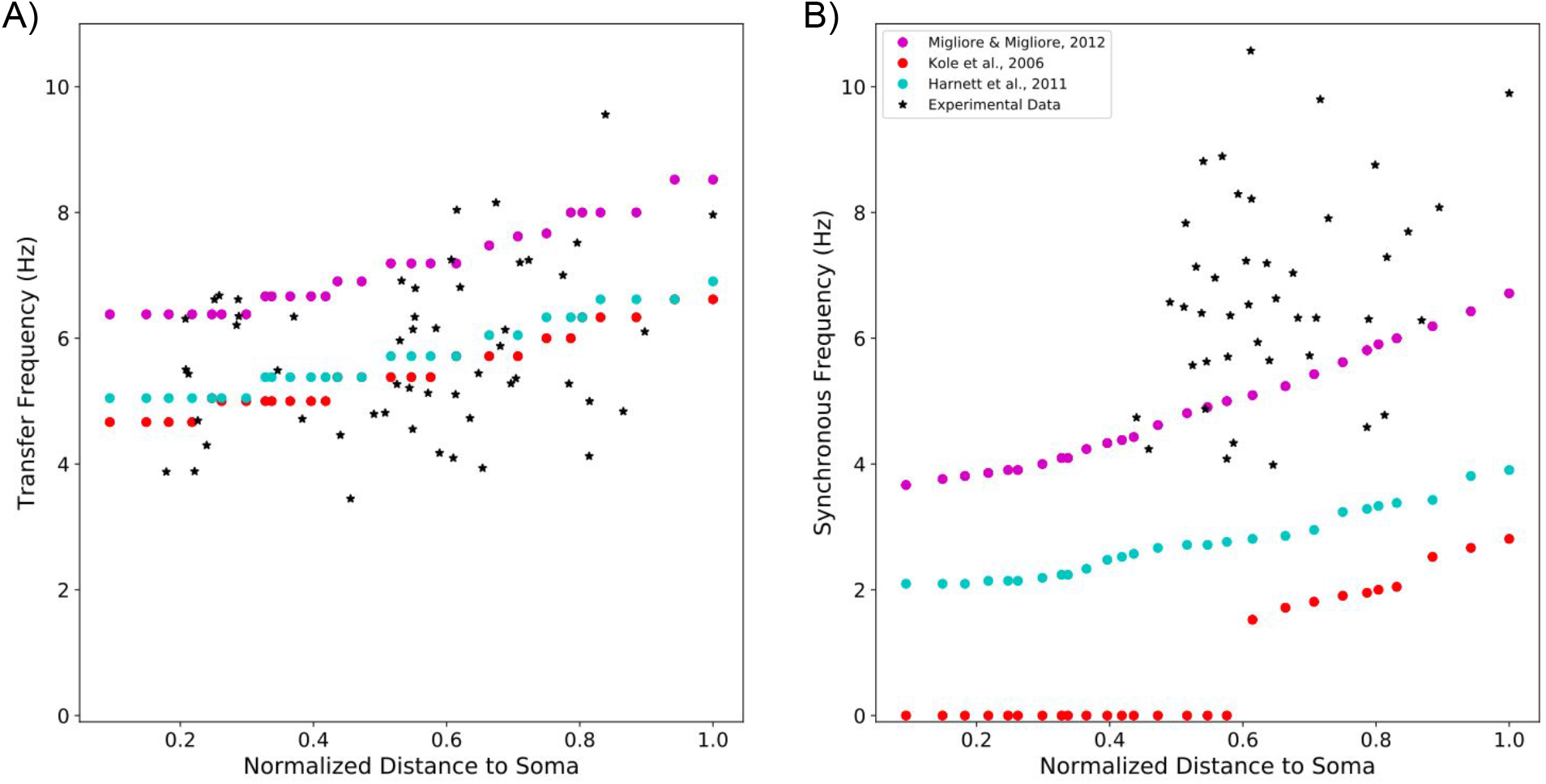
A model of HCN including a TASK-like shunting current best approximates experimentally observed impedance profiles. Resulting impedance features when using three different models of HCN channels in the same model neuron. (A) Compared to the original PT model which uses the HCN and TASK-like channel models from Migliore & Migliore (2012), the mechanisms developed by Kole et al. (2006) and Harnett et al. (2011) reduced transfer frequency along the apical trunk, but the values remain well within the experimental range. (B) They led to dramatic reductions in synchronous frequency however.

HCN mediates dendritic resonance and leading phase response in PTs, but TASK-like shunting current can modulate them (Fig. 6). By simulating the chirp stimulation along the apical trunk of **model 5** while blocking either *I*_h_ or the TASK-like shunting current across the entire neuron, we observed the independent effects of HCN and TASK-like channels on the impedance profile. Blocking *I*_h_ while leaving the shunting current intact increased impedance amplitude across frequencies but eliminated resonance and inductive phase, as expected from experiments [13, 30, 75]. Instead, both impedance amplitude and phase fell off with frequency as in a simple, passive parallel RC circuit model. Blocking the shunting current dramatically increased impedance amplitude, more so than blocking I_*h*_, but reduced transfer frequency, resonance strength, synchronous frequency, and impedance phase across frequencies (Fig. 6 A, B). It is noteworthy that blocking the shunting current did not reduce synchronous frequencies along the apical trunk to the level of the other two HCN models from Harnett et al. (2015) and Kole et al. (2006), but it did reduce them to the low end of the experimental range (Fig. 5). Therefore, the TASK-like shunting current alone does not endow a realistic phase response; the Migliore & Migliore’s model of *I*_h_ alone was sufficient for realistic synchronous frequencies. The changes to dendritic impedance caused by blocking I_*h*_ and shunting current were consistent along the apical trunk, becoming more pronounced at the distal end of the trunk where HCN and shunting current density were highest (Fig. 6C-F).

**Fig 6.**
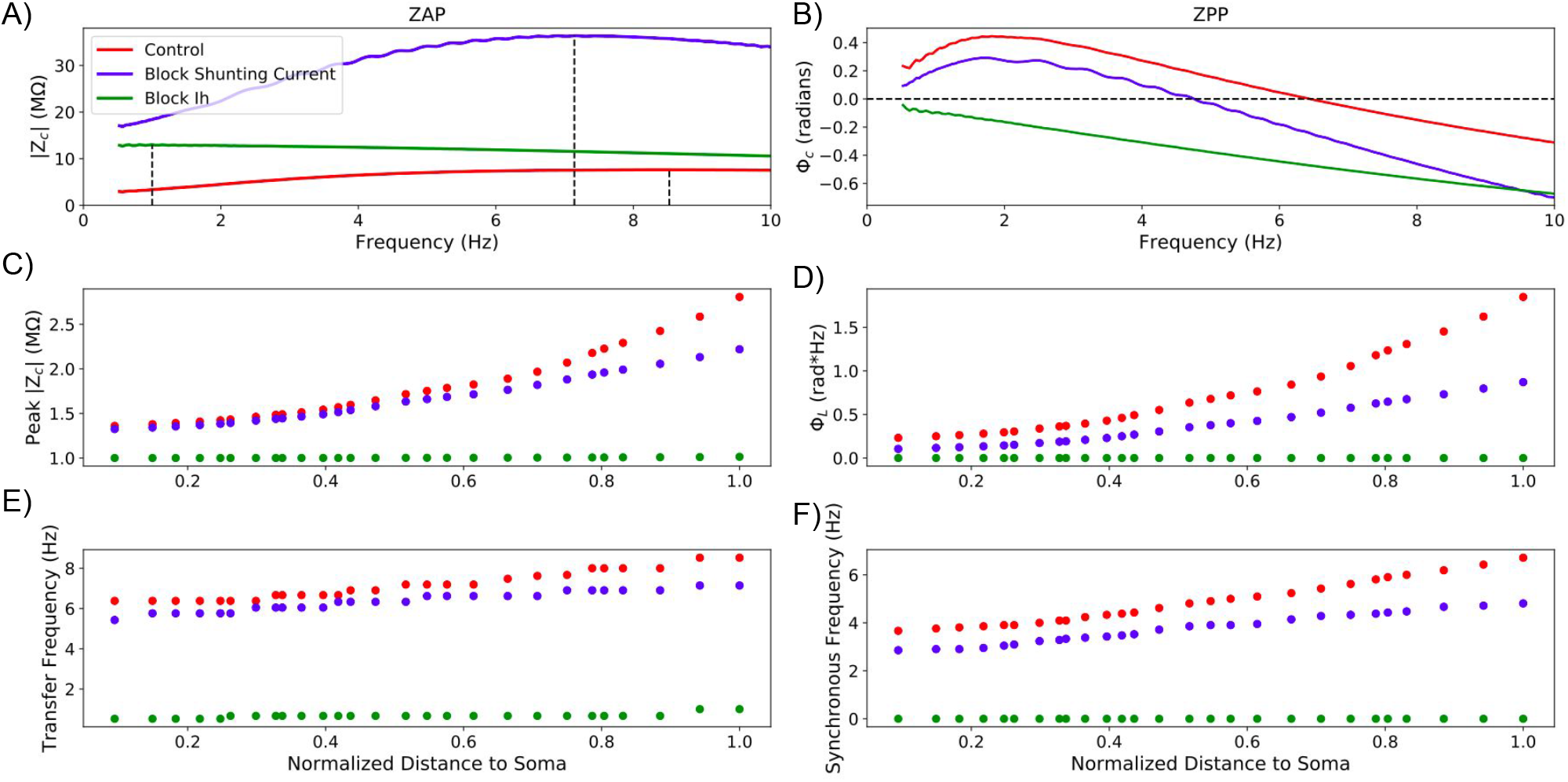
Selective blockade of *I*_*h*_ and shunting current differentially modulates dendritic impedance. Panels A & B show example ZAPs and ZPPs between the distal end of the apical trunk and the soma 288.9 *µ*m away, respectively, under baseline conditions (red) and when either I_*h*_ (green) or the shunting current (red) have been blocked. We also observe how (C) resonance strength, (D) total inductive phase, (E) transfer frequency, and (F) synchronous frequency are attenuated along the apical trunk under those same conditions.

Blocking HCN and TASK-like channels influences EPSPs in accordance with changes to dendritic impedance (Fig. 7). The downward shifts in ZPP caused by blocking HCN and TASK-like channels seen in Fig. 6D correspond to reductions in the compensation for membrane capacitance, increasing the delay in EPSP peak at the soma. Blocking TASK-like shunting current increases the lag between peak synaptic current halfway along the apical trunk and peak V_memb_ at the soma by 1 ms. Blocking I_*h*_ increases the lag by 2.6 ms. Similarly, blocking these currents reduces resonance strength (Fig. 6C) and the width of the EPSPs are increased accordingly. Although the changes to EPSP peak timing are fairly small, coupled with the changes in EPSP shape these channels can have a large impact on the integration and coincidence detection of synaptic potentials in the soma.

**Fig 7.**
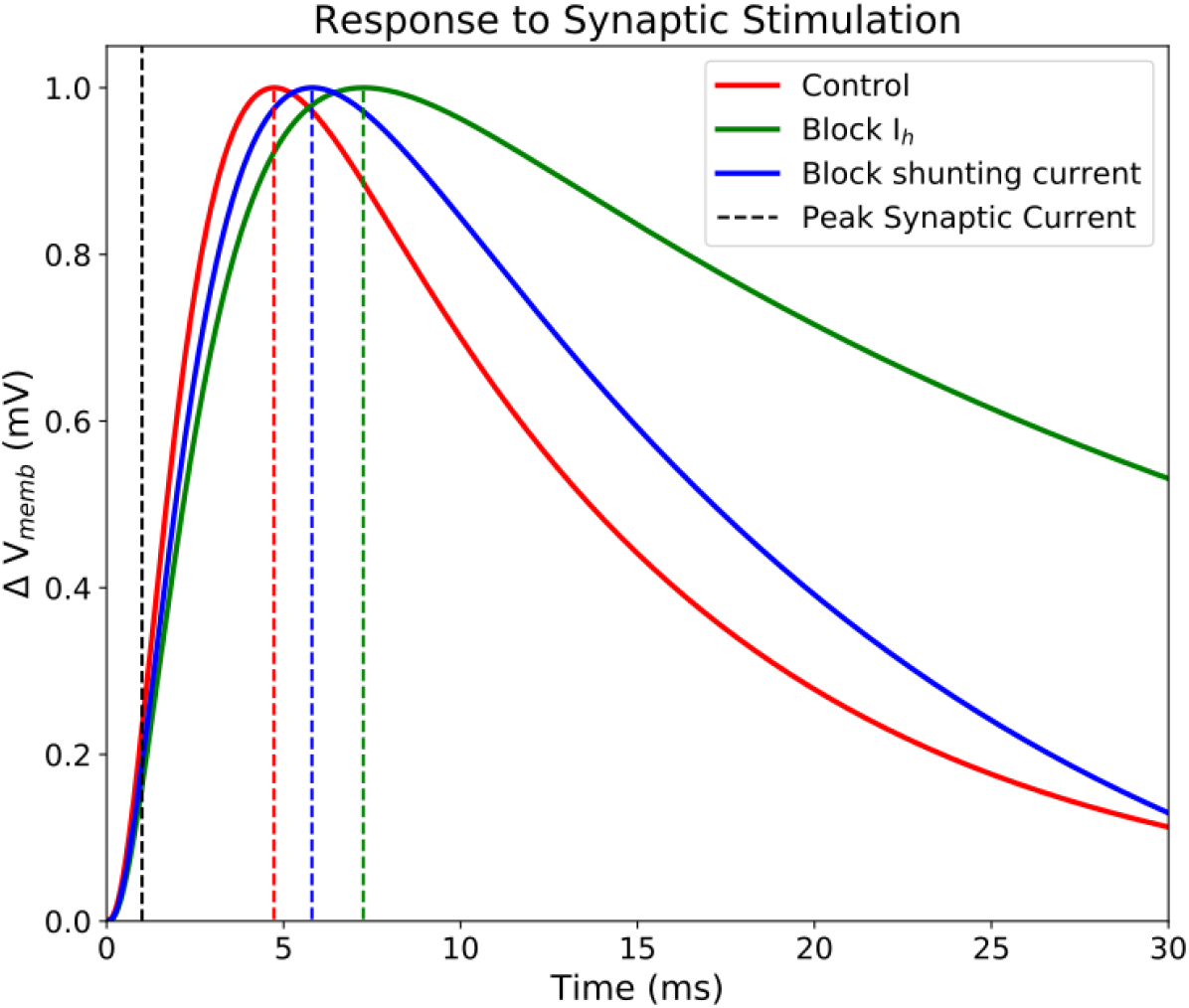
Selective blockade of I_*h*_ and shunting current effects timing and shape of EPSPs. **Model 5** was stimulated with an single excitatory synaptic current roughly halfway along the apical trunk (136.4 *µ*m from the soma), and V_memb_ was measured at the soma following blockade of HCN (green) and TASK-like channels (blue), as well as under control conditions (red). Maximal synaptic conductance was tuned to produce a 1 mV EPSP at the soma, and peak synaptic current occurred at 1 ms (black, vertical dashed line). Maximal EPSP V_memb_ lagged 3.7 ms behind peak synaptic under control conditions, 4.8 ms after blocking TASK-like shunting current, and 6.3 ms after blocking I_*h*_. EPSPs narrow in accordance with decreasing resonance strength seen in Fig. 6.

### Degeneracy and the interplay of I_*h*_ and shunting current

We demonstrated how different distributions of HCN and TASK-like channels can produce realistic impedance profiles (Fig. 8). This is an example of degeneracy, wherein different combinations of elements, parameters, or in this case ion channels can produce the same behavior. We replaced the HCN channel model used in **model 1** with the combined HCN and TASK-like channel models described by Migliore & Migliore (2012) and used in **model 5**. While we preserved the original HCN channel distribution from **model 1**, this replacement maintained realistic transfer frequencies (Fig. 8C) and produced synchronous frequencies which are in line with experimental observations (Fig. 8D). The location-dependent impedance response in the adjusted **model 1** was similar to that of **model 5**, even though the HCN and TASK-like channel distributions in **model 1** was exponentially increasing with distance from the soma across the full length of the apical dendrites, while in **model 5** their densities were constant in the apical tufts (Fig. 8B). This shows degeneracy of the impedance profile and is consistent with the variability of PTs seen *in vivo* [13, 75]. It also further demonstrates that the Migliore & Migliore (2012) implementation of I_*h*_ and TASK-like shunting current provides the most biologically plausible sources of inductive phase in neocortical PTs.

**Fig 8.**
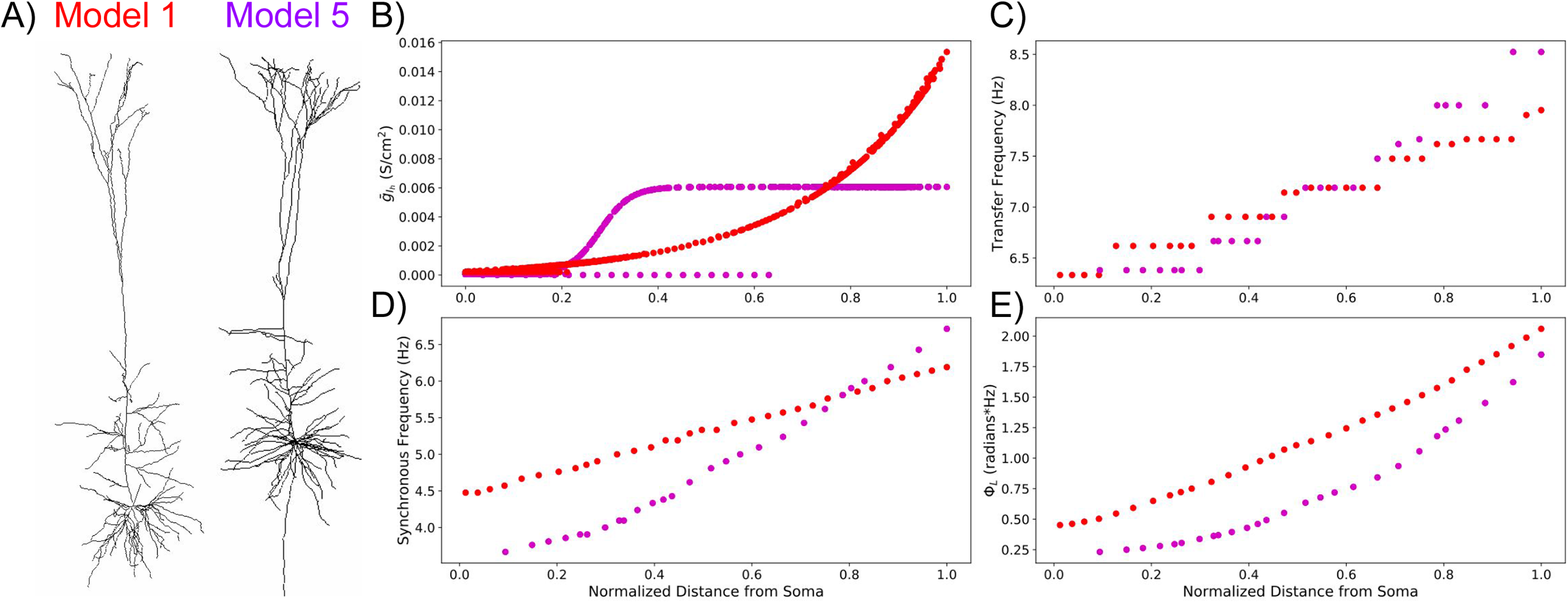
Using a model of HCN channels including a shunting current in model 1 produces realistic impedance amplitude and phase response, comparable to model 5. (A) Morphologies of **model 1** and **model 5**. (B) Distribution of 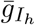, which affects both I_*h*_ and TASK-like shunting current, in the two models. Distances are normalized to the farthest compartment from the soma in each cell. (C) Transfer frequencies increase but remain within experimental range as originally. (D) Synchronous frequencies along the apical trunk are greatly improved compared to the experimental data. (E) Total inductive phase between **model 5** and the adjusted **model 1** are similar. Note that distances in C-E are normalized to length of each model’s apical trunk

Both I_*h*_ and TASK-like shunting current modulated all features of the impedance profile, but they did not contribute to the impedance profile equally (Fig. 9). By varying HCN and/or TASK-like channel density (ΔI_*h*_ and ΔI_*lk*_, respectively) by ±90% in increments of 10% uniformly across the dendritic arbor, we explored how different distributions of these channels in **model 5** modulated the dendritic impedance profiles. We measured how transfer impedance between the distal end of the apical trunk and the soma was affected by these changes to HCN and TASK-like channel densities (Fig. 9). Changes to HCN channel density had a greater impact on the impedance profile than equivalent changes to TASK-like channel density, but one can compensate for the other. For example, at baseline (ΔI_*h*_ = 0%; ΔI_*lk*_ = 0%), transfer frequency between the distal dendrite and the soma was 8.26 Hz; transfer resonance strength, 2.65; synchronous frequency, 6.32 Hz; and total inductive phase, 1.60 rad*Hz. By increasing HCN density across the neuron by 60%, transfer frequency increased to 8.97 Hz, resonance strength increased to 2.96, synchronous frequency increased to 7.06 Hz, and total inductive phase increased to 2.01 rad*Hz. These changes may be roughly compensated by also decreasing TASK-like channel density across the neuron by 70%, where transfer frequency was 8.26 Hz; resonance strength, 2.52; synchronous frequency, 6.42 Hz; and total inductive phase, 1.67 rad*Hz. Thus, regions of single color intensity in Fig. 9 represent degenerate combinations of HCN and TASK-like channel densities.

**Fig 9.**
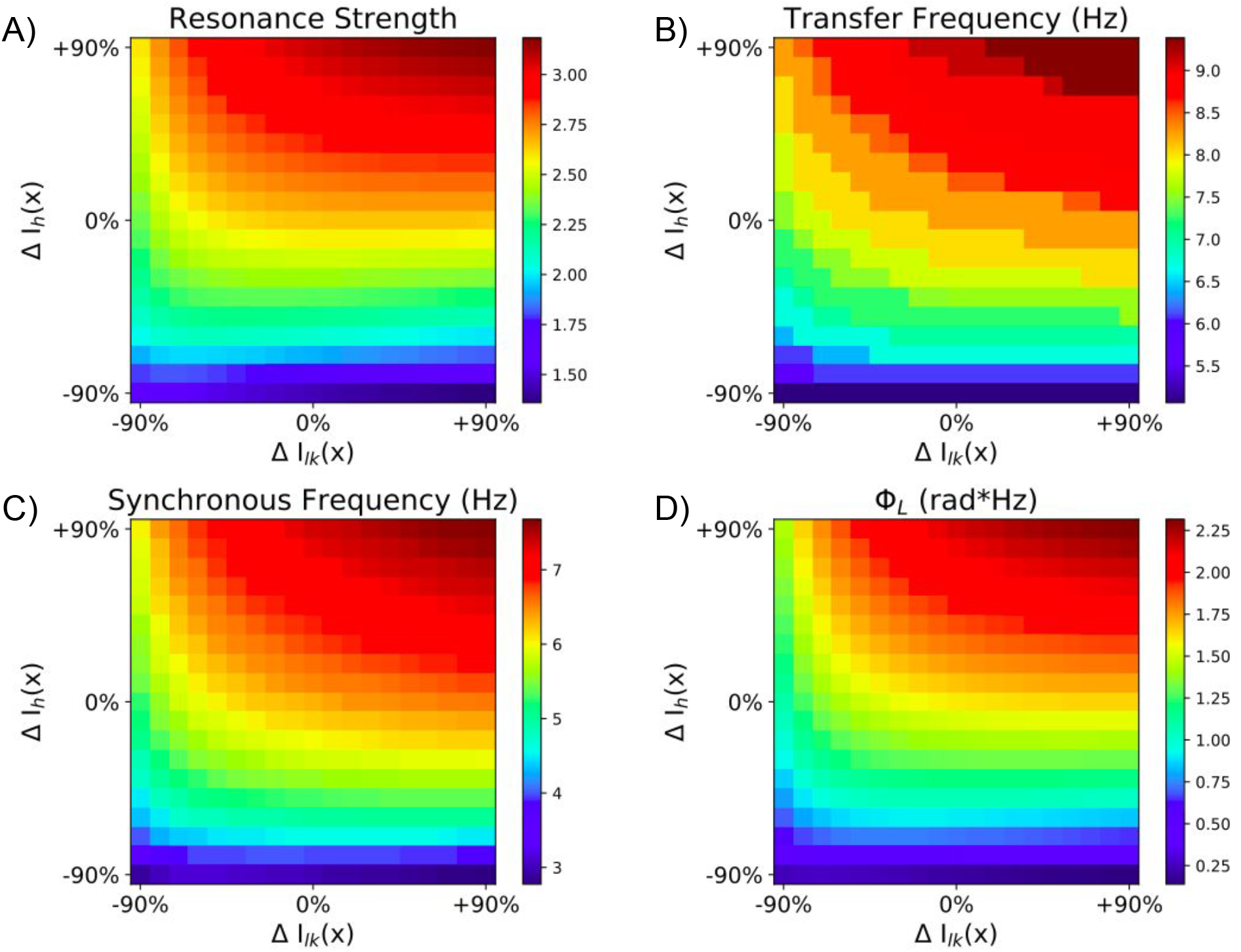
Combined effects of modulating HCN and TASK-like channel density on dendritic impedance. HCN density (ΔI_*h*_) and/or TASK-like channel density (ΔI_*lk*_) were modulated by ±90% in 10% increments across the entire neuron, which altered (A) resonance strength, (B) transfer frequency, (C) synchronous frequency, and (D) total inductive phase.

## Discussion

### Dendritic conductances and impedance profiles

While none of the models studied were explicitly designed to exhibit realistic dendritic impedance, only **model 5** accurately captured features from both impedance amplitude and phase profiles. The combination of I_*h*_ with a TASK-like shunting current, which mediated the realistic dendritic impedance we have seen, was intended to account for the paradoxical change from the excitatory to inhibitory effect of I_*h*_ in response to increasingly strong synaptic inputs [16, 19, 48]. The parameters of the shunting current were tuned to reproduce this result while maintaining an F-I curve consistent with experimental observations [16, 54]. Importantly, **model 5** did not reproduce biologically realistic phase response after eliminating the TASK-like channels and replacing the HCN model with those developed by Kole et al. (2006) and Harnett et al. (2015) (Fig. 5). With the other HCN channel models, **model 5** still maintained the experimentally observed F-I curve, but did not reproduce the paradoxical change from the excitatory to inhibitory effect of I_*h*_ in response to increasingly strong synaptic inputs. [20, 37, 54]. This suggests the model combining I_*h*_ and the TASK-like shunting current developed by Migliore & Migliore (2012) provides the best approximation of the currents mediating the location-dependent impedance profiles of PT cells.

It is also noteworthy that a combination of dendritic I_*h*_ and TASK-like shunting current undermined the hypothesis that the aforementioned paradoxical effect of I_*h*_ is mediated by M-type K^+^ channel currents [19]. Recent work has demonstrated the effects of M-type K^+^ channels on dendritic impedance in the lobula giant movement detector neurons in grasshoppers [14]. We do not, however, expect M-type K^+^ channels to have a significant impact on dendritic impedance in neocortical PTs. While the distribution of M-type K^+^ channels in neocortical PTs is poorly understood, they are rare in the dendrites of CA1 PCs [6]. The only model studied here that included M-type K^+^ channels throughout the dendrites was **model 3**, which produced reasonable synchronous frequencies but overestimated dendritic transfer frequencies. This can be attributed to **model 3’s** high HCN channel density, by far the highest of the five studied here (Table 1).

**Model 5** provides some insights into the dendritic ZPPs of PTs, with total inductive phase increasing by more than 150% along the length of its apical trunk, a far greater increase than seen in the other models (Fig. 4). Similar results were obtained in **model 1** after it was adjusted to include TASK-like channels. Though previous experimental studies have not described increases in total inductive phase with distance along the apical trunk in PTs, the relationships seen in **models 1 & 5** were comparable to those seen in CA1 PCs [52]. Leading or inductive phase is driven by the balance of membrane capacitance and phenomenological inductance and is therefore sensitive to the distribution of dendritic conductances [9, 12, 27, 28, 45, 46, 52, 66, 75, 76]. Our results support the notion that inductive phase, mediated by *I*_h_ and modulated by a TASK-like shunting current, provides a mechanism for compensating the location-dependent capacitive delay of dendritic inputs. This has been hypothesized as ensuring that simultaneous synaptic inputs distributed across the dendritic arbor are coincident at the soma [76]. Here we see that HCN channels and TASK-like shunting current, by contributing to inductive phase, both help to reduce the capacitive delay in the arrival of synaptic inputs to the soma (Fig. 7). Some have also suggested that inductive phase provides a mechanism by which subthreshold neuronal membrane oscillations might maintain phase relationships with ongoing local field potentials [12, 52, 76]. Although of the precise physiological role of inductive phase remains an open question, we believe a model with realistic dendritic phase response is more likely to have realistic distributions of dendritic ion-channels.

The resonance mediated by *I*_h_ in PTs qualitatively differs from resonance mediated by K^+^ currents seen in trigeminal root ganglion neurons from guinea pigs or photoreceptors in blowflies [24, 57, 57]. Resonant frequencies of the input ZAP at the soma in those cells range from 10 − 200 Hz, while resonant frequencies in PTs are in the range of 3 − 10 Hz [13, 29, 75]. Furthermore, both location-dependence and impedance phase remain largely unexplored in neurons with K^+^-mediated resonance.

Dendritic impedance profiles are not static. Previous work has demonstrated that subthreshold resonance can be dynamically tuned by ongoing activity [12, 27, 41, 51, 52, 61, 62, 70]. For instance, long-term potentiation induces changes in the impedance profile of hippocampal PCs [51]. Dynamic changes to impedance profiles may have a role in pathophysiology. For example, there is evidence for upregulation of HCN channel expression following epileptic seizure [42, 63, 69]. And while HCN channels are necessary for resonance in PTs, the dendritic impedance profile can be significantly altered upon modulation of other local conductances or morphological changes to the dendritic tree [15, 17, 28, 30, 32, 60, 62, 83]. A number of studies have explored the possibility of modulating the dendritic impedance profile by manipulating other channels like A- and M-type K^+^ channels or Ca^2^+ channel “hot-zones” [14, 17, 60]; however, the possibility of modulating dendritic impedance via TASK-like shunting current has not previously been investigated. Our observations of changes to the impedance features through changes to HCN and TASK-like channel density suggest a paradigm by which degeneracy and tunability of the impedance profile may arise (Fig. 9). For instance, changes to the impedance profile caused by changes in I_*h*_, either through changes in HCN channel expression or noradrenergic modulation [39], may be compensated for by appropriate adjustments to the TASK-like channel density, and vice versa . It is important to note, though, that changes to TASK-like shunting current cannot compensate for the complete absence of HCN channels in PTs, but can only modulate the impedance profile in its presence.

### Experimentally-verifiable predictions

1. **model 5** was the only unaltered model to reproduce experimentally observed transfer frequencies and synchronous frequencies, and it also exhibited much larger total inductive phase (Fig. 4A). Therefore, we expect this to be the case in real PTs as well. For instance, 2 Hz was the optimal leading phase between the soma and the center of the apical trunk 136.4 *µ*m away Fig. 4B). That 0.2 radian lead translates to a roughly 17 ms lead in peaks in somatic V_memb_ response compared to peaks in the current stimulus. While there is no doubt variability in PT dendritic impedance profiles, we expect a 17 ms lead is reasonable and probably at the lower end of the possible range.
2. If the shunting current’ is indeed produced by TASK channels, dendritic impedance should be reversibly alterable by changes to extracellular acidity, as TASK channels are pH sensitive [72].
3. The paradoxical effects of I_*h*_ observed by George et al. (2009) were abolished with application of the drug XE991. These effects are best accounted for by interaction between I_*h*_ and a TASK-like shunting current, and it has been suggested that XE991 may block TASK channels [16, 48]. Therefore, we predict that bath application of XE991 to PT cells will produce comparable changes to the dendritic impedance profile as those observed when blocking the shunting current in Fig. 6.
4. We also expect blocking the shunting current to produce an increased lag between peak synaptic current in the apical dendrites and peak somatic V_memb_, and blocking I_*h*_ should produce an even greater lag (Fig. 7).

### Limitations and future directions

A major limitation of this study is a limitation of most biophysically-detailed models of neurons: the distribution of conductances and passive properties are assumed to be either constant or vary smoothly along the neuronal topography, and this is often not the case [1, 49, 67]. Each of these properties can influence the dendritic impedance profile. For instance, hot-zones of Ca^2^+ channels have been shown to have an impact on dendritic impedance, but the precise parameters defining these hot-zones differed among the models presented here [16, 17, 23, 54]. Differences in the distribution of parameters illustrate the degeneracy of dendritic impedance, however. Both **model 5** and the adjusted version of **model 1**, which includes the TASK-like shunting current, similarly captured the features of the impedance profile observed along the apical trunk (Fig. 8). These two models had different morphologies, passive properties, Ca^2^+ channel hot-zone dimensions, and perhaps most importantly different distributions of HCN channels along the neurons’ longitudinal axes. Impedance analysis of the apical trunk could not, therefore, provide any indication as to whether either of these distribution schemes are more likely to exist in real neurons. Considering the degeneracy of dendritic impedance, these two distributions may be a small sample of a wide range of possible distributions. In fact, it is unlikely that diversity in morphology and passive properties alone can account for the variance of impedance features observed experimentally [13, 75]. Instead, these differences in the distribution of ion channels across the dendritic tree are likely strong contributors to the diversity of PT impedance profiles. Further investigation of the precise contributions of morphology and channel distribution to dendritic impedance in PTs is one avenue for our future work.

Our exploration of degeneracy of the impedance profile was also limited. We observed how uniform changes to HCN and TASK-like channel densities across the entire dendritic arbor produce different impedance profiles (Fig. 6 & Fig. 9). This would be analogous to cell-wide changes in channel expression, bath application of agonists or antagonists, or possibly changes to extracellular pH. We did not explore how localized changes to HCN and TASK-like channel activity may affect the impedance profile. For instance, stimulation of postsynaptic alpha2A adrenoceptors has been shown to inhibit HCN channel activity [79]. This is not to mention the influences of passive membrane properties, other active channels, or morphology on dendritic impedance [12, 31, 32]. Although showed how HCN channels and TASK-like channels may affect the impedance profile, determining how they interact with these other cell properties to modulate dendritic impedance profiles is a goal for future studies.

While we have focused on dendritic impedance in neocortical PTs from rodents, how these results relate to PTs in humans remains unclear. Recent studies have shown interesting differences between PTs in humans and rodents regarding I_*h*_-mediated phenomena like subthreshold resonance and sag potentials in the soma [5, 33]. Some of these results may be attributed to differing expression of HCN subtypes between species [33]. The I_*h*_-dependent physiological differences between rodents and humans are based on measurements from the soma, so how these results may extend to the dendrites is still an open question [5, 33]. Furthermore, the majority of data on resonance from human PTs comes from patients with epilepsy, which is associated with pathological effects on HCN channels [42, 63, 69] To better understand the relationship between the results presented here and dendritic impedance in human PTs, we need a better picture of the distribution of HCN and TASK-like channels in their dendrites and how they are affected in epilepsy. Considering the relative scarcity of human data compared to rodent data, and the difficulty of performing the experiments necessary for obtaining this information, computational modeling will be indispensable in bridging the gap between species.

## Acknowledgments

We wish to thank Michael Hines and Robert McDougal (Yale) for useful discussions on this subject. Supported by NIH R01EB022903, U01EB017695, R01MH086638; NSF Internet2 E-CAS 1904444; NYS DOH01-C32250GG-3450000; NIDCD R01DC012947-06A1; Army Research Office Grant W911NF-19-1-0402

## Abbreviations

PC: pyramidal cell
L5: layer 5
PT: pyramidal tract neuron
TASK channel: Twik-related acid-sensitive K+ channel
ZAP: impedance amplitude profile
ZPP: impedance phase profile
*I*_h_: h-current
HCN channel: hyperpolarization-activated cyclic nucleotide–gated
AMPA: *α*-amino-3-hydroxy-5-methyl-4-isoxazolepropionic acid
IT: intratelencephalic
CT: corticothalamic
EPSP: excitatory post-synaptic potential

